# A VLP-based immunogen that elicits selective anti-Myostatin antibodies, enhances muscle mass and strength, and reduces adiposity

**DOI:** 10.64898/2026.04.06.716693

**Authors:** Quiteria Jacquez, Julianne Peabody, Eduardo Hernandez Acosta, Bryce Chackerian, S. Joseph Endicott

**Affiliations:** Department of Pathology, University of New Mexico Health Sciences Center, Albuquerque, NM 87131, USA; Department of Molecular Genetics and Microbiology, University of New Mexico School of Medicine, Albuquerque, NM 87131, USA

**Keywords:** Myostatin, metabolism, aging, vaccine, virus-like particle, muscle, grip strength

## Abstract

Myostatin (MSTN) is a TGFβ family ligand that restricts muscle growth. Genetic loss-of-function in MSTN increases muscle mass, reduces fat accumulation, and improves metabolic health in mice and humans, with no known adverse phenotypes. Thus, depleting MSTN has therapeutic potential for obesity, sarcopenia, and other muscle wasting conditions. Recently developed monoclonal antibodies (mAbs) targeting MSTN or its receptors are expensive, require frequent injections/infusions, and risk a loss of efficacy from the development of anti-drug antibodies. Here, we report a comparatively inexpensive and durable alternative to mAbs, a virus-like particle (VLP)-based active immunotherapy, termed “MS2.87-97”, that elicits an antibody response against a discrete and unique epitope in mature MSTN protein, with no cross-reactivity to GDF11. Compared to controls, MS2.87-97-treated mice had less age-associated weight gain and exhibited significantly reduced body fat by DEXA scan. MS2.87-97-treated mice also had significantly improved bodyweight-adjusted grip strength, and upon dissection, they were found to have increased muscle mass. No major safety concerns were identified. Echocardiography revealed no evidence of functional impairment of the heart, and histological analysis showed no change in myocardial collagen deposition (fibrosis). These initial findings support the continued preclinical development of MS2.87-97 as an immunotherapeutic for treating obesity, sarcopenia, and muscle wasting.

## Introduction

Myostatin (MSTN) is a secreted ligand of the TGFβ family with both paracrine and endocrine signaling functions that inhibit satellite cell (muscle stem cell) activation^1^. During embryonic development, MSTN limits the number of muscle fibers that form, and in adults MSTN restricts muscle fiber size^2^. Humans with rare inherited loss of function mutations in MSTN are unusually muscular and strong, even as infants, and they show no major deleterious phenotypes^3^. People carrying a single nucleotide polymorphism in MSTN are disproportionally represented in some kinds of strength-based professional sports^4^.

Mice that are heterozygous knockouts for myostatin have enhanced muscle volume, increased strength, and live 15% longer than normal controls^5^. Homozygous knockouts have a normal lifespan^5^, but they show signs of improved healthspan in old age, such as improved cardiac function, reduced fibrosis in the myocardium, increased bone mineral density, and reduced circulating insulin and glucose^6^. MSTN null mice are resistant to fat gain and insulin resistance on a high-fat diet, and deletion of MSTN dramatically reduces adiposity and improves glucose tolerance in genetic models of obesity^7–9^. These findings suggest that blocking MSTN signaling could be highly beneficial for treating obesity/diabetes, sarcopenia, and conditions of muscle wasting, with a low risk of side effects.

MSTN is secreted as a homodimeric pro-protein, which is cleaved by the extracellular protease Furin, into the pro-domain and the mature domain, which remain non-covalently associated as latent MSTN^10^ until the pro-domain is cleaved by the Tolloid protease, releasing mature MSTN^11^. Mature MSTN signals primarily through the type II activin receptor ACVR2B, and to a lesser extent, ACVR2A^12^. Upon ligand binding, type II receptors engage the type I receptors ALK4 and ALK5, which are functionally redundant in MSTN signaling^13^. Activin A also signals through these receptor complexes, inhibiting muscle hypertrophy^14^. Wild-type mice injected with a soluble form of ACVR2B show an increase in muscle mass by 30-40% in two weeks, over PBS injected controls^14^.

Several pharmaceutical companies have developed humanized monoclonal antibodies (mAbs) to target multiple steps in the MSTN signaling pathway, including ACVRA/B (Bimagrumab)^15^, pro/latent MSTN (Apitegromab)^16^, mature MSTN (Trevogrumab)^17^ and Activin A/INHBA (Garetosmab)^18^. However, mAb-based therapies have the drawbacks of requiring frequent injections or infusions^19^, high cost, and a potential loss of efficacy over time due to the formation of anti-drug antibodies^20^.

We have developed a novel, bacterial-expressed, and scalable replacement for anti-MSTN mAbs – an immunotherapy that activates the immune system to produce selective anti-MSTN antibodies. This immunotherapy is based on highly immunogenic, non-infectious virus-like particles (VLPs), produced from the capsid protein of bacteriophage MS2, displaying an antigen unique to MSTN. Antigens displayed on highly dense, repetitive (multivalent) structures, such as VLPs, are immunogenic enough to efficiently activate anergic self-reactive B cells to produce antibodies^21,22^. Previous studies of this immunotherapy indicate that autoantigen-specific B cells internalize the VLP–antigen fusion and present foreign VLP-derived epitopes to T helper cells, resulting in T help without requiring MSTN-specific T-cell self-reactivity^23^. Although VLP-based display of self-antigens efficiently activates anti-self antibody responses, these immunogens do not elicit autoreactive T cells in mice^24,25^ or humans^26^.

We have found that an MS2-based VLP displaying an antigen representing amino acids 87-97 of the mature MSTN protein (termed “MS2.87-97”) elicits the production of antibodies with no cross-reactivity to GDF11 (the most similar protein to MSTN in the mouse and human proteomes). This VLP increases muscle mass, enhances grip strength and reduces adiposity. No deaths or other adverse events were observed in any of the mice, and assessments of heart health indicated normal cardiac function. Thus, MS2.87-97 is the first of a new generation of immunotherapies to harness the immune system to target MSTN and enhance metrics of musculoskeletal and metabolic health, without obvious side effects.

## Results

### Specificity: VLP immunogens are designed based on uniqueness to MSTN

*Mus musculus* mature MSTN is highly similar to other TGFβ family ligands (**Fig 1A**) and 89% identical to mature GDF11, an essential bone morphogenic protein. This presents a particular challenge in developing an immunotherapy that targets MSTN, but not GDF11. 100% of GDF11 knockout mice die within a few hours of birth^27^. Mice overexpressing follistatin (FST), a TGFβ ligand inhibitor that promotes degradation of both MSTN and GDF11, have increased muscle mass (from inhibition of MSTN), but they also have reduced bone density (from inhibition of GDF11), leading to spontaneous bone fractures^27^, underscoring the importance of selectively targeting MSTN over GDF11. Previous attempts to produce MSTN-based immunogens have all failed to target unique MSTN epitopes (**Fig 1B–C** )^28–33^. Targeting MSTN without also targeting GDF11 can only be accomplished with an immunization technology if it is capable of eliciting highly immunogenic responses against discrete unique epitopes. Unlike other vaccine platforms, VLPs are uniquely capable of producing strong antibody responses against short peptide epitopes^34^, ideal for discriminating MSTN from other TGFβ family ligands.

**Figure 1.**
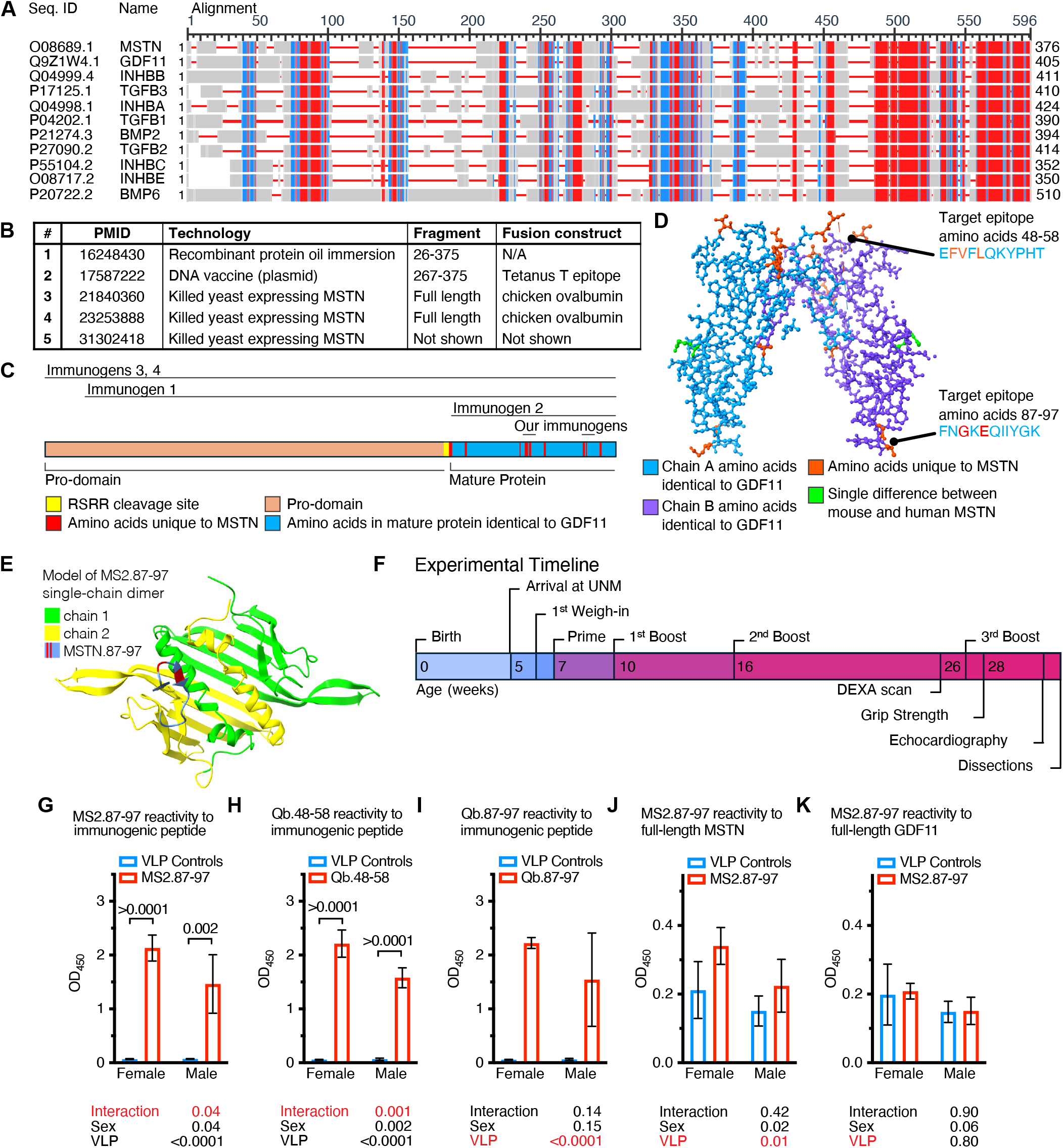
VLPs are designed to elicit antibodies against MSTN, with no reactivity to GDF11. (A). COBALT Sequence alignment of *Mus musculus* MSTN, and the ten most similar proteins in the mouse proteome (all TGFβ-family ligands). Red indicates high conservation, which is highest in the C-terminal part of the sequence that constitutes the mature protein. Blue indicates low conservation. Gray is non-conserved. (B) A table of previous studies that have attempted to make vaccines against MSTN in mice. (C) To-scale map of the primary structure of the *Mus musculus* MSTN protein, indicating the epitopes used by priory vaccines, and our new VLP-based immunogens. (D) Structure of the MSTN protein, colored by identity to GDF11 (blue and purple). Red amino acids are unique to MSTN over GDF11. The green amino acid indicates the single difference between the mouse and human orthologs. (E) The AlphaFold-predicted structure of the MS2.87-97 single chain dimer indicates that the MSTN epitope presents in a loosely ordered loop that projects from the outward facing surface of the MS2 coat protein. (F) The experimental timeline for evaluating the efficacy of MS2.87-97. (G–I) VLPs elicit strong serum reactivity, as determined by ELISA, against the peptides displayed by MS2.87-97, Qβ.48-58, and Qβ.87-97 VLPs, respectively, indicated high immunogenicity. (J) There was moderate serum reactivity of mice treated with MS2.87-97, against mature MSTN. (K) There was no detectable serum reactivity of mice treated with MS2.87-97, against mature GDF11. Error bars are standard deviations. The results of a full-model 2-way ANOVA are shown below the plots in G–K. In G&H, where there is a significant interaction of sex x VLP, the results of 2-tailed unpaired t tests are shown on the plots. For all experimental groups n = 4.

We selected two peptide sequences from the mature MSTN protein that contained differences from GDF11 (**Fig 1D**): amino acids 48-58 (EFVFLQKYPHT) or amino acids 87-97 (FNGKEQIIYGK). Four versions of the MSTN immunotherapeutic were synthesized for initial testing, using two separate virus-like particle (VLP)-based strategies: (1) synthetic MSTN peptides were chemically conjugated to Qβ bacteriophage VLPs^35,36^ and (2) MSTN epitopes were cloned into the surface-exposed loops of the single chain dimer of the MS2 bacteriophage capsid protein, as described previously^37^. Of the two MS2-based designs, only the design presenting the MSTN 87-97 epitope (termed “MS2.87-97”) was both readily soluble in aqueous solution and also capable of forming VLPs. The AlphaFold-predicted structure of a single MS2.87-97 protein is shown in **Fig 1E**.

Four male and four female BALB/cByJ mice were immunized with the three different VLPs at age 7 weeks (by intramuscular injection without exogenous adjuvant), and then boosted at ages 10, 16, and 27 weeks, as shown in the experimental timeline (**Fig 1F**). Three weeks after the first boost, all three VLP vaccines elicited strong antibody responses against their respective displayed peptides (**Fig 1G–I**), but only MS2.87-97 elicited antibodies that cross-reacted with full-length MSTN. At age 32 weeks, when the animals were dissected, serum from MS2.87-97-treated animals showed modest immunogenicity against mature MSTN protein, with no detectable immunogenicity against GDF11 (**Fig 1J–K**). The subsequent sections will only report results from MS2.87-97 treated mice, since the other two VLPs failed to elicit immunogenic responses against the full-length MSTN protein and failed to induce physiological changes in the animals (not shown). As controls, groups of mice received unmodified (wild-type) MS2 or Qβ VLPs. The negative control groups showed no significant differences from each other and (in subsequent sections) were combined into a single VLP negative control group.

### Efficacy: MS2.87-97 increases muscle mass and reduces adiposity

Mouse body weights were measured twice weekly, prior to and following the first injection with VLPs. MS2.87-97-treated animals gained less weight over time, compared to negative controls that received unmodified VLPs (**Fig 2A–B**). A repeated measures 2-way ANOVA indicated a highly significant interaction effect of VLP x time, for both females and males. In order to understand how the reduction in body weight affected mouse body composition, we used dual energy X-ray absorptiometry (DEXA) scans to estimate fat content, lean content, and bone mineral content in anesthetized mice at age 26 weeks (approximately 4.5 months after the first injection). Qualitatively, the mice treated with MS2.87-97 appeared thinner than the mice treated with undecorated VLPs (**Fig 2C**). The DEXA scan indicated that mice treated with MS2.87-97 had reduced total fat mass (**Fig 2D**), reduced fat mass as a percentage of total body mass (**Fig 2E**), and reduced total body mass (**Fig 2F**). We did not detect a difference in bone mineral content (**Fig 2G**). Thus, a reduction in fat mass appears to be a major contributor to the reduction body weight in MS2.87-97-treated animals.

**Figure 2.**
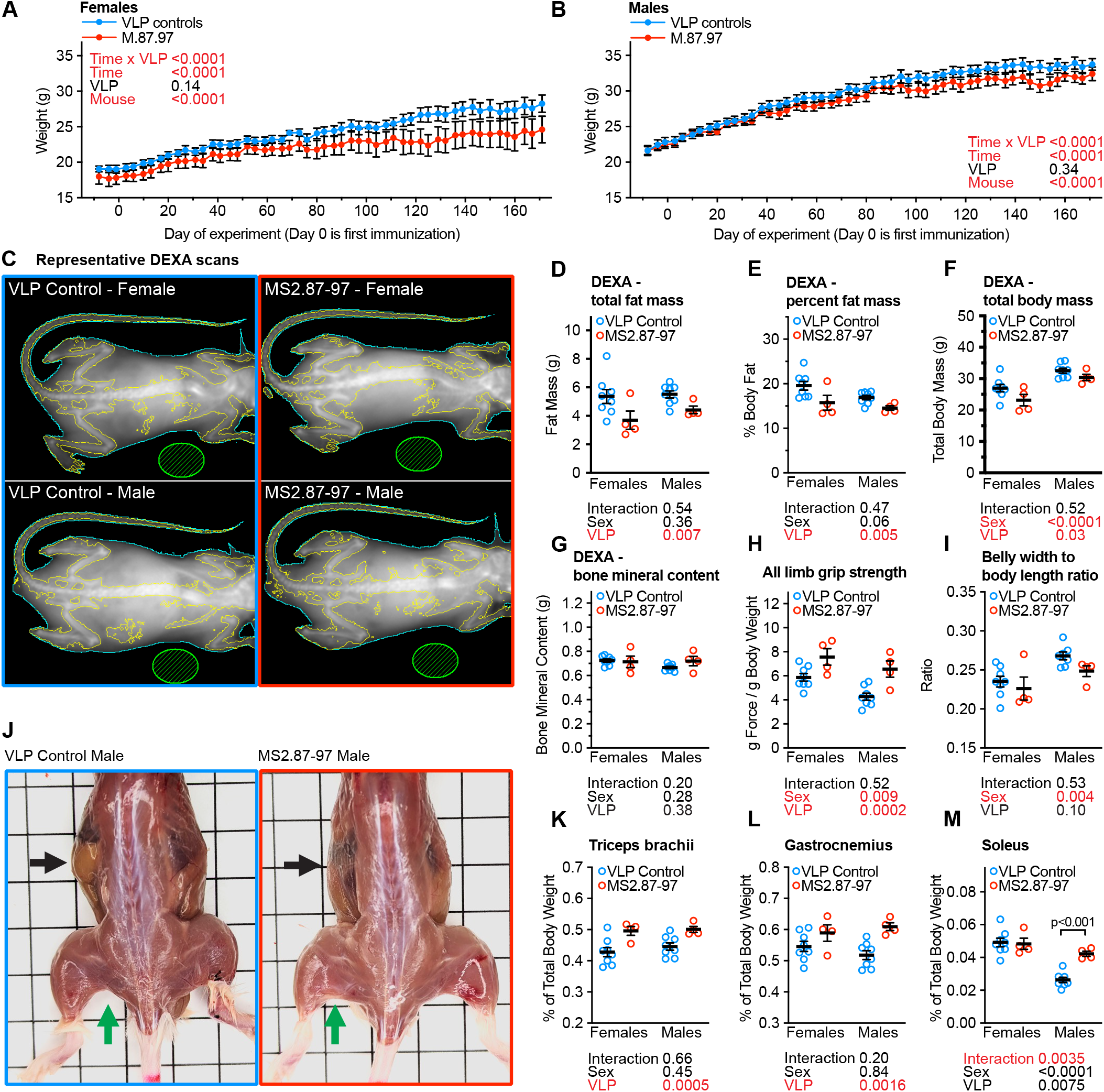
MS2.87-97 reduces adiposity and increases muscle mass and strength. (A). Weights of MS2.87-97 and VLP control treated female mice, across the experimental timeline. (B). Weights of MS2.87-97 and VLP control treated male mice, across the experimental timeline. (C). Representative DEXA scans of MS2.87-97 and control VLP-treated mice. (D) DEXA-estimated total fat mass of MS2.87-97 and control VLP-treated mice. (E) DEXA-estimated percent of body mass made of fat from MS2.87-97 and control VLP-treated mice. (F) DEXA-estimated total body mass of MS2.87-97 and control VLP-treated mice. (G) DEXA-estimated bone mineral content of MS2.87-97 and control VLP-treated mice. (H) All-limb grip strength, as measured by a grip strength meter, of all experimental mice. (I) Ratio of maximum abdominal width to body length (nose to base of tail), measured post-mortem. (J) Representative images of skinned male mice from the MS.87-97 treatment group and the control VLP treatment group. Black arrows indicate the external oblique muscles, over the approximate position of the stomach. Green arrows indicate the position of the biceps femoris. (K) The weight of the triceps brachii muscle groups. (L) The weight of the gastrocnemius muscle (M) The weight of the soleus muscle. In K–M, each data point represents the average weight of both muscles (or muscle groups) from one animal. Error bars are S.E.M. The results of a repeated measures 2-way ANOVA are shown on the plots in A & B. The results of a full-model 2-way ANOVA are shown below the plots in D–M. In M, where there is a significant interaction of sex x VLP, the results of 2-tailed unpaired t tests are shown on the plot. For all experiments, MS2.87-97 n = 4 of each sex. Control VLP-treated animals were pooled for n = 8 of each sex.

We next sought to evaluate whether MS2.87-97 had an effect on muscle function. At age 28 weeks, we performed grip strength tests, and found that MS2.87-97-treated mice had significantly higher bodyweight-adjusted grip strength (**Fig 2H**). At age 32 weeks, the animals were humanely euthanized, measured, skinned, photographed on a 1 cm grid, and dissected. From the photographs, abdominal width (at the widest point) was determined, and plotted as a ratio to body length (nose tip to base of tail). There was a trend toward a reduction in this ratio, which did not reach statistical significance (**Fig 2I**). We tested for outliers by the IQR method (interquartile range), we found one outlier in the female MS2.87-97 treated group. If the 2-way ANOVA is run without this outlier, then the VLP effect is statistically significant (p = 0.007). One of the control males was excluded from this photographic analysis, because of a mistake that prevented the acquisition of a high-resolution photograph.

The photographs were examined for qualitative differences. MS2.87-97-treated animals had a thickening of the external oblique muscles, resulting in an opaque lateral body wall with visible striations (**Fig 2J**). While the stomach (black arrow in **Fig 2J**), liver, and intestine can be clearly seen through the translucent body wall of the control animals, the thickened and opaque body wall of the MS2.87-97-treated animals obscures this view. The gluteus maximus and biceps femoris (green arrow in **Fig 2J**) were visibly enlarged. To quantitatively examine for changes in muscle mass, the triceps brachii, gastrocnemius, and soleus muscles were dissected and weighed. The triceps brachii (**Fig 2K**) and gastrocnemius (**Fig 2L**) were significantly larger (as a percentage of total body mass) in MS2.87-97-treated animals. For the soleus, MS2.87-97 treatment increased the muscle mass only in males (**Fig 2M**).

### Safety: MS2.87-97 did not have observable adverse effects on heart health

Only one study of MSTN depletion in mice has ever reported an adverse effect: inducible knockout of MSTN in the adult myocardium is cardiotoxic and lethal in male C57BL/6J mice^38^. This finding is controversial, and other papers have attributed the cardiotoxicity to the use of lethal doses of tamoxifen to induce the MSTN knockout^39,40^. There were no premature deaths in any the MS2.87-97 or control treatment groups. However, as a precautionary measure, we examined the hearts of MS2.87-97 mice for signs of damage.

At age 31 weeks, echocardiograms were performed. Female mice are less responsive to volatile anesthetics than male mice, and they recover much faster from cessation or interruptions in anesthetic administration^41^. Therefore, in our hands, collection of echocardiography data from males was more straightforward and reliable, and we are only reporting echocardiograms from male mice. We did not find differences in the diameters of the left ventricles, systolic or diastolic, (**Fig 3A-B**). Traces across the long axis of the left ventricle (**Fig 3C**) were used to assess ejection fraction (**Fig 3D**) and fractional shortening (**Fig 3E**), and no significant differences were detected.

**Figure 3.**
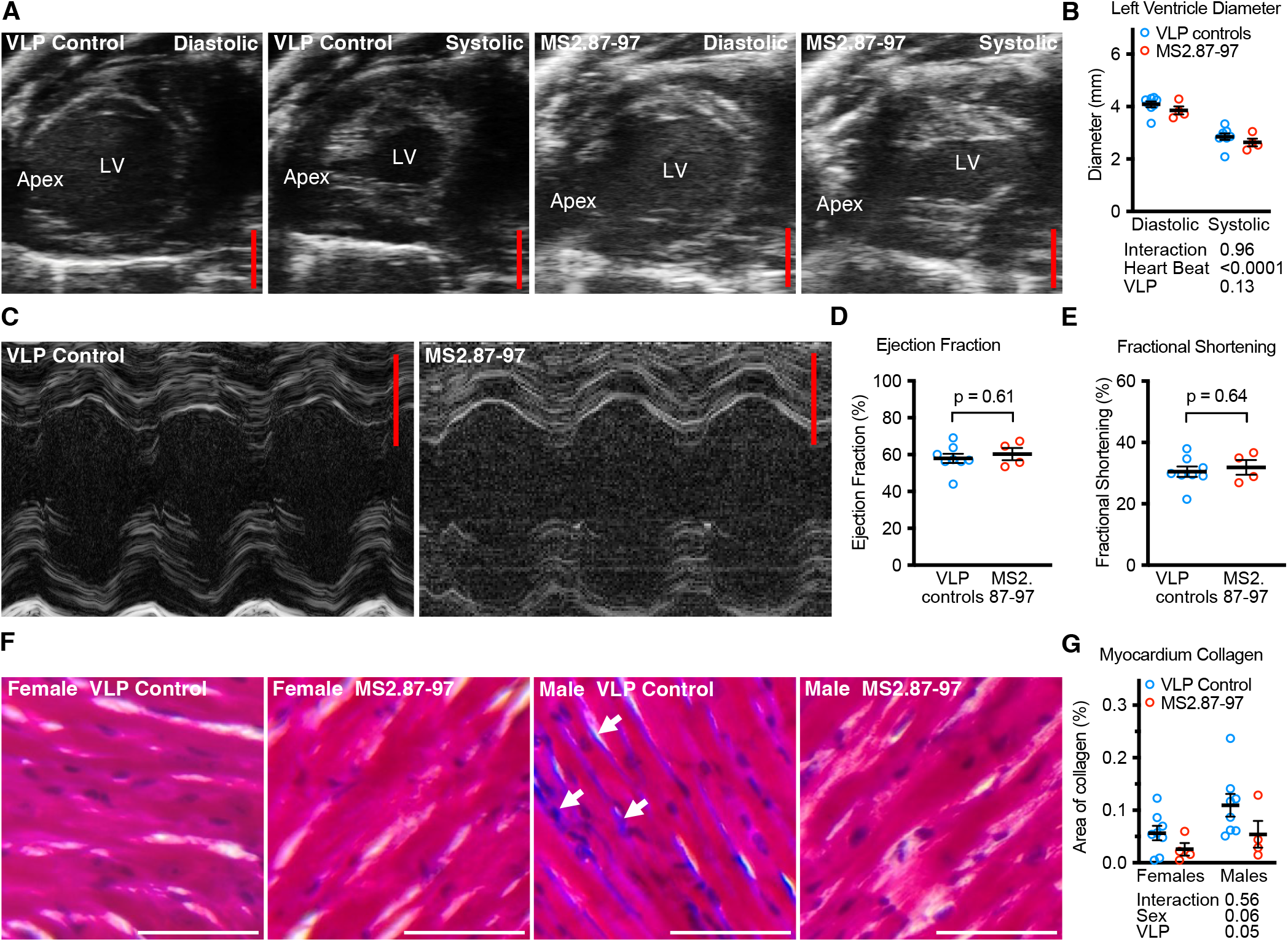
MS2.87-97 treatment does not produce observable effects on cardiac muscle function. (A) Representative echocardiograms of the left ventricles from male mice treated with either control VLPs or MS2.87-97. Scale bar = 2 mm. (B) Quantifications of the maximum (diastolic) and minimum (systolic) diameters of the left ventricle. (C) Representative M-mode images through the left ventricle from male mice treated with either control VLPs or MS2.87-97. Scale bar = 2 mm. (D) Ejection fractions from hearts shown in I. (E) Fractional shortening from hearts shown in I. (F) Representative images of transverse sections of the left-ventricular myocardium from females and males treated with VLP control or MS2.87-97, stained with Masson’s trichrome, showing muscle fibers in red, nuclei in dark purple, and collagen in bright blue. (G) Quantifications of the percentage of area in the myocardium that are positive for collagen staining (blue). White arrows in (F) indicate small amounts of collagen staining in the myocardium from a control male. The results of a 2-way ANOVA are shown below the plots in B, D, E, & G. Error bars shown in D & E are from an unpaired t test. For all experiments, MS2.87-97 n = 4 of each sex. Control VLP-treated animals were pooled for n = 8 of each sex.

Collagen deposition in the heart (fibrosis) has previously been used as a proxy for heart aging (where more collagen staining indicates a more aged heart) in MSTN deficient mice^6^. It was previously found that myocardial collagen deposition is no different between young adult (4-5 month) MSTN null mice and controls. However, at very old age (27-30 month), MSTN null mice have significantly less collagen deposition than controls, albeit still more than the young mice^6^. To examine for fibrosis in the hearts of MS2.87-97 treated mice and controls, transverse sections of ventricular myocardium were stained with Masson’s trichrome (**Fig 3F**). The percent of heart area that was positive for collagen staining trended to be less in MS2.87-97 treated mice, compared to controls, but this fell short of statistical significance (p = 0.05; **Fig 3G**). Collectively, these results indicate no signs of impaired functionality of the hearts in mice treated with MS2.87-97.

## Discussion

Here, we report the development and initial preclinical evaluation of a VLP-based active immunotherapy that elicits an antibody response that is selective for MSTN, over the closely related essential bone morphogenic protein GDF11. MS2.87-97 treated mice gained less weight with age and exhibited reduced adiposity by DEXA analysis. Skeletal muscle mass was enhanced, and grip strength was increased. Echocardiography revealed no evidence of heart functional impairment, and histological analysis showed a trend toward a decrease in myocardial collagen deposition. Together, these findings demonstrate that active immunization against a discrete MSTN epitope can promote favorable changes in musculoskeletal and metabolic health in mice.

While the initial results of this study are promising, there were some important limitations. First, as part of a pilot study, these experiments used a small sample size of isogenic (inbred to homozygosity) mice. Future studies will leverage larger sample numbers of outcrossed mouse stocks, in order to determine the generalizability of the findings. Second, MS2.87-97 elicited only modest immunogenicity against mature MSTN. However, previous studies have shown that even slight reductions in MSTN are sufficient to induce significant physiological changes. For example, mice that are heterozygous for the *MSTN* gene have only a 30% drop in circulating MSTN protein, but they show increases in muscle force generation of the same magnitude as *MSTN* knockouts^5^. While the lifespan of MSTN knockout mice is indistinguishable from wildtype controls, *MSTN* heterozygous mice live 15% longer. Thus, it appears that optimal MSTN dosing for lifespan and healthspan is a reduction that is somewhere between the wildtype and the knockout levels.

Previous studies have reported conflicting findings pertaining to the relationship between MSTN and cardiac health. A study of control and MSTN KO C57BL/6J mice at ages 27-30 months (older than median lifespan) revealed improved fractional shortening, slightly smaller left ventricular diameter (both systolic and diastolic), and reduced cardiac fibrosis^6^. These findings indicate that mice with MSTN knockout from birth have healthy hearts, which are perhaps even healthier than those in wildtype littermates. However, when a tamoxifen-inducible system was used to delete MSTN in the myocardium of young adult male C57BL/6J mice, it led to ventricular hypertrophy and 26.5% lethality within 10 days^38^. These findings have been disputed, because the dose of tamoxifen that was used has been found to be cardiotoxic by other labs^39,40^. Our analysis of heart health did not reveal any signs of MS2.87-97 cardiotoxicity. MS87-97-treated mice trended toward reduced cardiac fibrosis (which was just shy of statistical significance, at p = 0.05), consistent with the finding that aged knockouts have reduced cardiac fibrosis^6^. It might be the case that we did not detect a significant difference because we sacrificed the animals at middle age, before high levels of cardiac fibrosis are readily detectable.

MSTN and INHBA (Activin A) both inhibit muscle hypertrophy, and simultaneously blocking both MSTN and INHBA has a combinatorial effect, dramatically improving muscle growth^42^. However, Activin A is induced in cardiomyocytes under stress conditions, and it protects cardiomyocytes from ischemic injury^43^. Therefore, there is reason to believe that neutralizing Activin A could pose a risk of mortality in humans with risk factors for heart attack. Consistent with these concerns, in a recent clinical trial testing a combination of monoclonal antibodies targeting MSTN (Trevogrumab) and INHBA (Garetosmab) for their ability to prevent muscle wasting in humans taking semaglutide, it was found that the combination preserved more muscle mass than Trevogrumab alone. However, two patients receiving Garetosmab died of cardiac-related events, while no one died in the other study arms (see clinical trial NCT06299098). Similarly, in a clinical trial using Garetosmab as a treatment for fibrodysplasia ossificans progressiva, five participants in the Garetosmab treatment group died, but there were no deaths in the other study arms^44^. Thus, our finding that MS2.87-97 is safe for the heart underscores the value in targeting only MSTN, while leaving the remainder of TGFβ family ligands untouched.

There are a number of potential therapeutic applications for inhibiting MSTN, with the most obvious being the treatment of obesity. According to the CDC, over 40% of American adults are now obese. Common complications of obesity include type 2 diabetes mellitus^45^, hypertension, stroke, coronary artery disease, sleep apnea, non-alcoholic fatty liver disease, and some kinds of cancer^46^. Extreme obesity can shorten lifespan by as much as 14 years^47^. Glucagon-like peptide-1 receptor agonists (GLP-1 RAs) have emerged as effective treatments for weight loss by suppressing food consumption^48^, and one out of every eight (12%) of American adults has taken (or are currently taking) GLP-1 RAs to help with weight loss. However, ∼40% of weight loss induced by semaglutide (Ozempic) is lean mass (mostly skeletal muscle)^49^, and ∼25% of weight loss on tirzepatide (Zepbound) is lean mass^50^.

Adjunctive therapies, such as infusions of monoclonal antibodies that inhibit MSTN pathway signaling enhance fat loss and preserve muscle mass in mice^19^ and humans treated with GLP-1 RAs (see clinical trials NCT05616013 and NCT06445075). However, monoclonal antibody-based therapies have the drawbacks of requiring frequent injections/infusions^19^, high cost, and the potential loss of efficacy over time due to the formation of anti-drug antibodies^20^. MS2.87-97 is specifically designed to overcome these limitations as a comparatively inexpensive and scalable replacement for anti-MSTN monoclonal antibodies. If successfully brought to the clinic as an anti-obesity treatment, MS2.87-97 (or next generation anti-MSTN immunotherapies) could inexpensively lower non-communicable disease risk and give back years of healthy living to a hundred million Americans.

VLP-display of self-antigens has been successfully used in animal models to target self-molecules that are involved in the pathogenesis of diverse chronic diseases mediated by self-antigens, including Alzheimer’s Disease, hypertension, inflammatory diseases, and certain cancers^51^. Seven bacteriophage VLP-based immunotherapies have advanced into Phase I or II human clinical trials^52^, including vaccines for Alzheimer’s Disease (targeting amyloid-beta)^53^, allergy (targeting dust mite allergen)^54^, and hypertension (targeting angiotensin II)^55^. These vaccines have been shown to be well-tolerated and highly immunogenic. These highly promising findings suggest that the platform technology (VLPs) has a pathway for regulatory approval, pointing to the feasibility of the future clinical development of MS2.87-97 (or next generation anti-MSTN immunizations).

## Materials & Methods

### VLP production and purification

To produce MS2.87-97, primers containing the sequence encoding the MSTN epitope (amino acids 87-97) were used to generate a PCR product, which was then cloned into the downstream surface-exposed AB-loop of the MS2 coat protein single-chain dimer, utilizing BamHI and SalI restriction sites in the pDSP62 MS2 VLP expression plasmid^37,56^. The plasmid was amplified in Lucigen *E. coli* 10G ELITE Electrocompetent Cells (Fisher Scientific: NC0749147) and purified by miniprep (Qiagen: 27104), and successful cloning was confirmed by sequencing (Azenta).

The MS2.87-97 encoding plasmid was transformed into C41 (DE3) (Lucigen: 60341-1) *E. coli* cells by electroporation. Transformed C41 cells were grown at 37 °C in LB containing 50 µg/mL kanamycin until the cells reached an OD_600_ of 0.9, and protein expression was induced using 0.4 mM isopropyl-β-D-1-thiogalactopyranoside (Sigma: I6758-1G), and grown overnight at room temperature.

Cell pellets were collected by centrifugation, re-suspended in lysis buffer [50 mM Tris-HCL, 100 mM NaCl, 10 mM ethylenediaminetetraacetic acid, pH 7.5] with 1mg/ml hen egg lysozyme, and incubated on ice for 1 hour. Deoxycholate was added to 0.05% (w/v). The cell lysate was subjected to 3 rounds of sonication at 20 Hz for 30 seconds each. Cellular DNA was digested using 1mg/ml DNase I, 2.5mM MgCl_2_ and 0.5mM CaCl_2_ incubated at 37 °C for 1 hour. The lysate was then clarified by centrifugation (10,000xg, 10 minutes, 4°C), and the supernatant was purified by size exclusion chromatography (SEC) using a Sepharose CL-4B column (Cytiva: 17015001) pre-equilibrated with Sepharose column buffer (40 mM Tris-HCl, 400 mM NaCl, 8.2 mM MgSO4, pH 7.4). Fractions containing VLPs were identified by agarose gel electrophoresis. VLPs were concentrated from SEC purified fractions by precipitation with (NH_4_)_2_SO_4_ to 70% saturation and centrifugation followed by resuspension and dialysis in PBS (pH 7.4). The final concentration and purity of VLPs was determined via SDS-PAGE using a 10% NuPAGE gel (Invitrogen: NP0301BOX) using known concentrations of hen egg lysozyme as standards.

### Experimental Animals

All animal experiments were approved by the UNM HSC Animal Care and Use Committee (IACUC), protocol number 25-201572-HSC.

Five-week-old BALB/cByJ male and female mice were purchased from the Jackson Laboratory, and were housed in a specific pathogen free (SPF) room in the UNM HSC Animal Resource Facility (ARF). The animals were housed four to a cage, and each cage was randomly assigned to a treatment group. At 7, 10, 16, and 27 weeks of age, 5 μg of VLPs were administered by intramuscular injection.

### Blood Draws and Immunogenicity Assays

For live blood collections, mice were sedated using 3% isoflurane (Piramal: NDC66794-013-25) with oxygen as a carrier for 3 min, and approximately 50-100 μL of blood were collected by retroorbital bleed. Blood was allowed to coagulate at room temperature, and sera were separated by centrifugation at 1000 x g for 10 minutes.

At the conclusion of each study, mice were anesthetized with 4-5% isoflurane with oxygen as a carrier until they were unresponsive to toe pinch. Then the mice were exsanguinated by terminal cardiac puncture, collecting approximately 1 mL of whole blood per animal. Sera were prepared by the same methods, as above.

Antibody responses were quantified by ELISA using recombinant full length MSTN (PeproTech: 120-00-10UG). Immulon 2 ELISA plates (Thermo Scientific: 14-245-61) were coated with 50ng of recombinant MSTN (PeproTech: 120-00-10UG) in 50μL of PBS per well overnight at 4°C. Wells were blocked with 100 µL PBS-0.5% nonfat dry milk for one hour at room temperature. Sera isolated from immunized mice were diluted in PBS-0.5% milk and applied to wells in a 50μl volume for 2.5 hours at room temperature or overnight at 4°C. HRP-labeled goat anti-mouse IgG was diluted 1:5000 in PBS-0.5% milk (Jackson Immunoresearch: 115-035-003) and applied to wells in a 50μL volume for one hour at room temperature. Bound antibodies were detected by addition of 50μL TMB substrate (ThermoFisher: N301), and reactions were stopped using 50μL 1% HCl. Optical density was measured at 450nm using a Synergy HTX multi-mode plate reader.

### Dual-energy X-ray absorptiometry (DEXA)

DEXA scans were performed using the Lunar PIXImus Densitometer located in the Animal Resource Facility at UNM HSC, on mice anaesthetized by isoflurane. The Lunar PIXImus software was used to automatically calculate lean mass, fat mass, bone mineral content, and bone mineral density from the DEXA images.

### Echocardiography

Mice were sedated with isoflurane, and sedation was maintained by isoflurane/O_2_ by nose cone for the remainder of the procedure. Fur on the anterior chest was be depilated with Nair, and the skin was cleaned. Ultrasound was performed with the Visual Sonics Vevo3100 system and Vevo LAZR-X software, which is equipped with a temperature-controlled mouse pad that contains four-electrocardiogram (ECG) leads position under each limb. A rectal thermometer was used to monitor body temperatures throughout the procedure. The transducer was placed on the left hemithorax using pre-warmed aquasonic gel. Images were initially acquired in a parasternal long axis view. Short axis views were obtained subsequently. Using the 2D parasternal long-axis imaging plane as a guide, an LV M-mode tracing will be obtained close to the papillary muscle level. Vevo Lab software cardiac package was used to analyze 3 images each.

### Histology

Gastrocnemius, soleus, and heart were fixed in 3.7% formalin for 24 hours. The Human Tissue Repository and Tissue Analysis Shared Resource performed paraffin embedding and transverse sectioning to a thickness of 10 μm. Transverse sections of the heart, halfway through the length of the ventricles, were deparaffinized, rehydrated, and stained with Masson’s Trichrome (AbCam: ab150686). Collagen deposition in the myocardium was performed using the color thresholding tool in ImageJ, with blue color staining calibrated to images of the pericardium (positive control for blue collagen staining).

### Sequence alignment

The constraint-based multiple alignment tool (COBALT) version 1.26.0 was used to align the protein (amino acid) sequence of pro-MSTN and the pro-forms of the10 most similar proteins in the mouse proteome, using the default color scheme (conservation). Red indicates high conservation, blue is low conservation, and gray represents parts that are not conserved.

### Protein structure display

The protein structure display items were generated with Chimera X version 17. The crystal structure of mature MSTN^57^ is from the Protein Data Bank ID# 5JI1. The structure of MS2.87-97 was predicted using the AlphaFold Server (v3).

## Data Availability Statement

Data will be made available upon reasonable request to the corresponding authors.

## Acknowledgments

This project was supported by a pilot grant from the UNM HSC Research Allocations Committee. This project utilized services provided by core facilities and shared resources supported by P20GM121176 (UNM HSC Autophagy, Inflammation, and Metabolism Center) and P30CA118100 (UNM HSC Comprehensive Cancer Center).

## Author Contributions

Author contributions, according to applicable roles in the Contributor Role Taxonomy are: Conceptualization (SJE, BC), Data curation (QJ, JP, SJE), Formal analysis (SJE, QJ, JP), Funding acquisition (SJE, BC), Investigation (QJ, JP, EHA), Resources (BC, JP, SJE), Validation (JP, QJ), Visualization (SJE, QJ), Writing – original draft (SJE), Writing – review & editing (all authors).

## List of Figure Legends

